# De novo masking domains that gate TNF-family ligand assembly and activity

**DOI:** 10.64898/2026.04.20.719557

**Authors:** Shota Kudo, Brian Kuhlman

## Abstract

Tumor necrosis factor family ligands (TNFLs) are central regulators of immunity and promising agents for cancer therapy, but their clinical use is often limited by dose-limiting systemic toxicity. Conditional activation and genetic fusion with antibodies could improve safety and pharmacokinetics, yet these features are difficult to combine because TNFLs form obligate homotrimers that are structurally mismatched with antibody architectures. Here we use AI-enabled protein design to create de novo protein masks that conditionally control TNFL assembly. The masks are genetically fused to TNFLs through protease-sensitive linkers and prevent trimer formation by competitively binding the TNFL trimerization interface. We demonstrate with TNFα, OX40L, and 4-1BBL that protease-mediated release of the mask promotes trimer assembly, receptor binding and biological activity. These monomeric, switchable TNFLs are readily incorporated into full-length IgG formats, enabling plug-and-play construction of conditionally activatable antibody fusions. These fusions can be designed either to release activated soluble TNFLs upon protease treatment or, by reordering the fusion architecture, to trigger antibody multimerization and signaling by membrane-type TNFLs. When this multimerization strategy is applied to antibody-drug conjugates, conditional trimerization enhances cell killing by ∼30-fold, thereby improving the therapeutic window and enabling multiple strategies for selective tumor microenvironment targeting.

## Introduction

Cytokines are central regulators of immune responses and have long been recognized as promising therapeutic agents for cancer treatment. Indeed, several cytokines have been clinically approved for oncology indications^1, 2^. However, their broader clinical application has been limited by a narrow therapeutic window, primarily resulting from rapid systemic clearance, pleiotropic biological activities, and dose-limiting toxicities^3^. Over the past decades, extensive protein engineering efforts have been undertaken to address these challenges, including antibody or antibody fragment–mediated targeting strategies, mutational optimization of receptor binding affinities and specificities, cytokine co-localization approaches, and prodrug formats designed to enable localized activation^4, 5, 6^.

Tumor Necrosis Factor Ligands (TNFLs) comprise a family of 19 cytokines that regulate survival, differentiation, and functional responses of both immune and non-immune cells^7, 8^. Most TNFLs are type II membrane proteins, some of which are released as soluble factors upon proteolytic processing. Functionally, TNFLs exist as obligate homotrimers and engage Tumor Necrosis Factor Receptors (TNFRs) in a defined 3:3 stoichiometry, in which receptor clustering and higher-order organization are essential for productive signaling^9, 10^. Several TNFLs, including TNFα, OX40L, and 4-1BBL, have emerged as attractive candidates for cancer immunotherapy due to their potent activation of innate and adaptive immune responses^11, 12^.

TNFα, for example, has been approved in Europe for the treatment of soft tissue sarcoma. However, its clinical use remains severely constrained by systemic toxicity. To improve pharmacokinetic properties and achieve localized activity, antibody fusion strategies and prodrug approaches have been developed. Notably, immunocytokines consisting of an scFv targeting the extra-domain B of fibronectin fused to TNFα demonstrate preferential tumor accumulation^13, 14^, while TNFα prodrugs, which are fusion proteins that mask the receptor-binding surface of the TNFα trimer and incorporate linkers cleavable by tumor-associated proteases^15, 16^, show reduced systemic toxicity and localized biological activity^17, 18^. AcTakine, consisting of a low-affinity TNFα mutein fused to a targeting VHH, improves safety while maintaining in vivo efficacy^19^. Several of these strategies have progressed into clinical evaluation.

Despite these advances, the safe and effective delivery of TNFLs remains a major challenge. An optimal delivery format would combine prolonged systemic exposure with strict conditional activation at the target site. However, the development of such formats is inherently complicated by the structural mismatch between TNFLs, which function as trimers, and antibodies or Fc domains, which are intrinsically dimeric. This incompatibility has limited the design of stable fusion architectures capable of simultaneously achieving long half-life and conditional cytokine activation.

We hypothesized that if we could create a monomeric form of TNFL that was stable and soluble but then could be conditionally proteolyzed and released to form active trimer, then we would have a molecule that would be well-behaved as an antibody fusion. This format would allow tumor targeting via the antibody followed by local activation of the TNFL with tumor-enriched proteases. In considering the discovery of binding/masking proteins that stabilize the TNFL monomer, one important consideration is that traditional experimental approaches for binder discovery, such as animal immunization (antibody discovery) or yeast display (general protein binders), are not possible since wild type TNFL does not present as a monomer, it is an obligate homotrimer. In contrast, computational modeling enables the virtual dissociation of TNFL trimers, thereby exposing otherwise inaccessible interfaces and rendering them amenable to rational design. Recent advances in AI-based protein design^20, 21^ have made it possible to systematically explore such non-native structural states, providing a framework to disrupt native oligomerization while preserving fold stability.

In this study, we develop a stable and generalizable platform for the safe delivery of TNFLs by rationally targeting their obligate trimeric architecture. Leveraging AI-driven protein design, we engineer a novel masking domain that disrupts TNFL homotrimerization and stabilizes artificial monomeric TNFL variants by engaging the native trimerization interface. This computational framework integrates generative structural design^22^, sequence realization under structural constraints^23^, and independent structure validation^24^, enabling access to protein states that are difficult to achieve experimentally. We first establish proof of concept using TNFα and subsequently demonstrate the versatility of this approach by extending it to OX40L and 4-1BBL. Finally, we introduce two antibody fusion formats that leverage these monomeric TNFL designs to enable conditional delivery of both soluble and membrane-type TNFLs. Beyond cytokine therapeutics, these antibody architectures also provide a foundation for the development of conditionally active antibody–drug conjugates (ADC) with an expanded therapeutic window.

## Results

### Design of masking domains for human TNFα

We designed masking (inhibitory) domains that target the native trimerization interface of human TNFα using an AI-based protein design strategy (Fig. 1a). An artificial monomeric TNFα structure was generated from the crystal structure of the TNFα homotrimer (PDB ID: 1A8M) by computationally separating the individual protomers. Using this monomeric representation, we identified hydrophobic hotspots within the native trimerization interface, which are expected to contribute to trimer stability and thus represent attractive targets for disruption. We then generated 1,000 backbone scaffolds for masking domains against these sites using RFdiffusion. After filtering based on radius of gyration, 182 backbone models were retained. For each backbone, 10 sequences were designed using ProteinMPNN, yielding a total of 1,820 candidate masking domain sequences.

**Figure 1.**
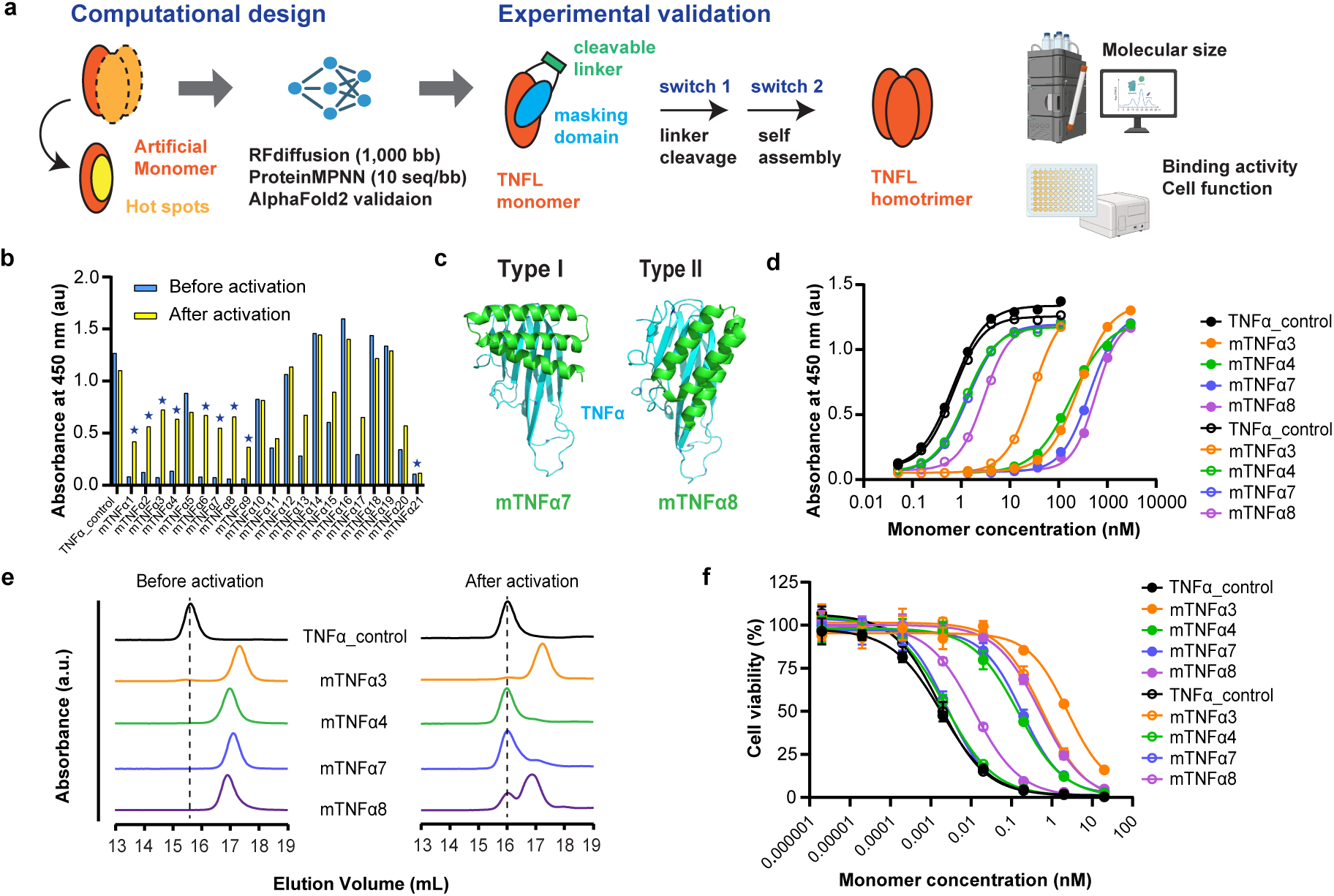
Computational design, screening and characterization of masked TNFα. A de novo masking domain was designed to bind the TNFα trimerization interface and stabilize an inactive monomeric state, enabling protease-triggered reassembly and recovery of TNFα function. Unless otherwise noted, protein concentrations are reported per TNFα protomer. a, Overview of the computational and experimental workflow used to design and prioritize masking domains for human TNFα. b, Initial ELISA screen using plate-immobilized adalimumab as a surrogate readout for TNFα trimer assembly. Adalimumab recognizes an epitope spanning two TNFα protomers. Signals before (blue) and after TEV activation (yellow) are shown. Asterisks indicate hit clones with reduced binding before activation. c, AlphaFold2-predicted complex structures of representative hit clones (mTNFα7 and mTNFα8). TNFα is shown in cyan and the masking domain in green. d, TNFR2 binding of selected masked TNFα constructs measured by ELISA before (filled circles) and after (open circles) TEV activation. Data are from a single experiment and are shown as representative results. e, Oligomeric states of selected masked TNFα constructs assessed by analytical SEC before (left) and after (right) TEV activation. The dashed line indicates the elution volume of the trimeric TNFα control. f, Cytotoxic activity of selected masked TNFα constructs in WEHI-13VAR cells before (filled circles) and after (open circles) TEV activation. Cell viability (%) was calculated as 100 × (S − B)/(V − B), where S is the sample signal, V is the 100% viability control (cells plus medium), and B is the 0% viability control (cells plus 1% Triton X-100). Data are shown as mean ± s.d. from n = 3 technical replicates.

All candidates were subsequently evaluated using AlphaFold2 (AF2) as an independent structure prediction framework to assess global fold stability and preservation of the intended interface geometry. Of the 1,820 designs, 119 met stringent structural criteria, including high model confidence (pLDDT > 85), robust interface prediction (iPTM > 0.85), and minimal deviation between backbone conformations before and after sequence design (RMSD < 1.5 Å). From these, 21 designs exhibiting diverse backbone geometries were selected for experimental characterization. The sequences of the selected candidates, along with key AF2 metrics, are summarized in Supplementary Data and Supplementary Table 1.

### Screening and characterization of masked TNFα

Designed masking domains and the TNFα monomer were expressed as fusion proteins, with a TEV protease–cleavable linker (GSGGGGSGENLYFQSGGGGS) inserted between the two components to enable efficient screening of masking designs. As a trimeric control, wild-type TNFα carrying the same TEV-cleavable linker at the N-terminus was also expressed. Proteins were produced in Expi293F cells in a 96-well format, and expression levels were evaluated by SDS–PAGE (Supplementary Fig. 1a). Expression in the culture supernatant was comparable across constructs, with typical yields of ∼0.3 mg per mL of culture.

For the initial screen, we used adalimumab binding as a surrogate readout for TNFα assembly. The crystal structure of the TNFα–adalimumab Fab complex (PDB ID: 3WD5) shows that adalimumab recognizes a composite epitope spanning two adjacent TNFα protomers within the homotrimer and overlaps with the TNFR-binding site. Thus, enforced monomerization is expected to reduce binding, whereas trimer reassembly should restore the epitope and increase apparent binding under surface-immobilized conditions.

Adalimumab was immobilized on ELISA plates, and culture supernatants were analyzed either before or after TEV treatment. Bound proteins were detected using an anti-FLAG–HRP conjugate. Before activation, the trimeric TNFα control gave a strong signal, whereas 9 of 21 fusion proteins showed substantially weaker binding (Fig. 1b). After TEV treatment, the control remained strongly positive, and 8 of these 9 low-signal constructs, except mTNFα21, showed a marked increase in signal, consistent with recovery of a higher-order TNFα arrangement recognizable by adalimumab after linker cleavage.

AlphaFold predictions for these 8 responsive constructs suggest that all masking domains adopt a three-helix-bundle fold and bind at the native TNFα trimerization interface (Supplementary Fig. 1b). Two distinct binding modes were observed (Type I and Type II). In Type I designs, exemplified by mTNFα7, the helical bundle is oriented approximately perpendicular to the TNFα β-sheet surface, whereas in Type II designs, exemplified by mTNFα8, the helical bundle is oriented more parallel to the β-sheet surface (Fig. 1c).

Based on backbone diversity and predicted binding modes, we selected four constructs (mTNFα3, mTNFα4, mTNFα7, and mTNFα8) for further characterization. These proteins were expressed in Expi293F cells and purified by IMAC followed by SEC. The main peak was collected and used for subsequent analyses (Supplementary Fig. 1c).

We first assessed binding to TNFR2 by ELISA (Fig. 1d). TNFR2 was immobilized on plates, and binding was measured before and after TEV activation. The trimeric TNFα control showed similar binding with or without TEV treatment, whereas the fusion proteins exhibited substantially stronger binding after activation, consistent with the screening results. EC50 values varied across constructs, and mTNFα4, mTNFα7, and mTNFα8 showed >100-fold stronger apparent binding after activation than before activation (Supplementary Table 2). A similar activation-dependent increase was observed to adalimumab (Supplementary Fig. 1d, Supplementary Table 3), and TEV cleavage under these assay conditions was near-complete (Supplementary Fig. 1e).

We next evaluated TNFα homotrimer reassembly after activation using analytical SEC (Fig. 1e). Prior to activation, each fusion protein eluted as a single peak, with later retention volumes (16.9–17.3 mL) than the trimeric TNFα control (15.6 mL), consistent with a smaller hydrodynamic species. After TEV treatment, the trimeric TNFα control eluted as a single peak at

16.0 mL. In contrast, all fusion proteins resolved into two peaks: an earlier peak at 16.0–16.1 mL and a later peak at 16.9–17.2 mL. Because the earlier peak co-eluted with the trimeric control, we assigned this species as trimeric TNFα, whereas the later peak was consistent with the monomeric fusion species observed prior to activation. TEV cleavage was similarly efficient across constructs under these conditions (Supplementary Fig. 1f). Together, these data indicate that TEV cleavage drives TNFα homotrimer reassembly. Importantly, the abundance of the earlier, higher-order peak correlated with ELISA binding strength, consistent with avidity arising from multimerization.

Finally, we evaluated biological activity using the murine WEHI-13VAR sarcoma cell line, which is highly sensitive to TNFα and is widely used for assessing the function^25^. The trimeric TNFα control exhibited comparable potency with or without TEV treatment. In contrast, prior to activation, all fusion proteins showed >75-fold reduced activity relative to the control (Fig. 1f, Supplementary Table 4). Following TEV activation, biological activity increased, although the extent of recovery differed substantially among constructs. Consistent with the ELISA and SEC analyses, mTNFα7 showed the largest shift in EC50 (>70-fold) upon activation. Comparable TEV cleavage efficiency was observed across constructs (Supplementary Fig. 1g).

Collectively, these results support our hypothesis that enforced monomerization of TNFα markedly reduces receptor binding and biological activity, and that TEV-mediated linker cleavage enables recovery of function through homotrimer reassembly accompanied by dissociation of the masking domain from the TNFα monomer.

### Extension of the masking strategy to OX40L and 4-1BBL

To evaluate the generalizability of our TNFL masking strategy, we extended the design framework to additional human TNFLs, namely OX40L and 4-1BBL, and performed computational design and prioritization of corresponding masking domains.

We explored three backbone-design strategies for the masking domains (Fig. 2a). Method 1 used standard RFdiffusion, as in the TNFα campaign, to generate 500 de novo backbones. Method 2 employed scaffold-guided RFdiffusion using the masking domains of TNFα (mTNFα7 and mTNFα8) as templates, generating 100 backbone variants. Method 3 used partial diffusion starting from an artificial complex between a TNFα masking domain and human 4-1BBL (PDB ID: 6BWV), generating 100 backbone variants. For this approach, a 4-1BBL monomer was superimposed onto the TNFα monomer in the mTNFα7 or mTNFα8 complex to transfer the interface geometry before diffusion. Methods 1 and 2 were applied to OX40L, whereas all three methods were applied to 4-1BBL.

**Figure 2.**
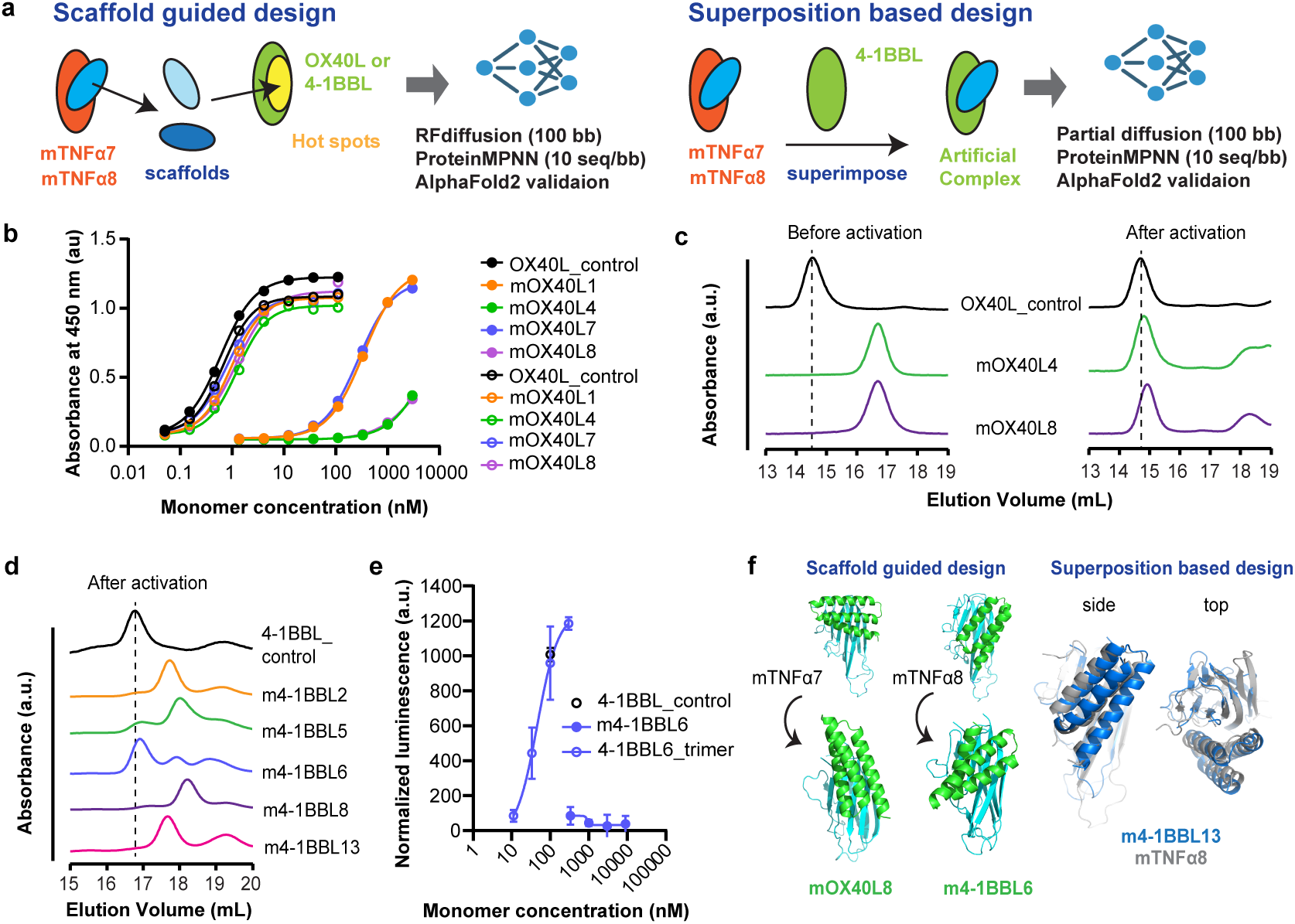
Design, screening and characterization of masked OX40L and 4-1BBL. To evaluate the generality of the masked TNFL platform, we extended the design and screening workflow to the additional TNFL family members OX40L and 4-1BBL using expanded computational design strategies. Unless otherwise noted, protein concentrations are reported per monomeric protomer. a, Expanded computational strategies for masking-domain design. In the scaffold-guided approach, the masking domains from mTNFα7 and mTNFα8 were used as structural scaffolds to guide backbone generation against the trimerization interfaces of OX40L and 4-1BBL. In the superposition-based approach, artificial complexes were generated by superimposing TNFα and 4-1BBL monomers, followed by partial diffusion to generate new 4-1BBL masking-domain backbones using the TNFα masking domains as templates. b, OX40 binding of selected masked OX40L constructs measured by ELISA before (filled circles) and after (open circles) TEV activation. Data are from a single experiment and are shown as representative results. c, Oligomeric states of selected masked OX40L constructs assessed by analytical SEC before (left) and after (right) TEV activation. The dashed line indicates the elution volume of the trimeric OX40L control. d, Oligomeric states of selected masked 4-1BBL constructs assessed by analytical SEC after TEV activation. The dashed line indicates the elution volume of the trimeric 4-1BBL control. e, Functional activity of masked 4-1BBL (m4-1BBL6) assessed in Jurkat-Lucia h4-1BB reporter cells. Responses to the monomer (filled circles) and SEC-purified homotrimer (open circles) are shown. TEV-treated 4-1BBL_control was included as a positive control. Data are shown as mean ± s.d. from n = 3 technical replicates. f, AlphaFold2-predicted complex structures of representative masked OX40L and masked 4-1BBL constructs, highlighting designs from the scaffold-guided and superposition-based approaches.

For each designed backbone, we generated 10 sequences using ProteinMPNN and screened designs with AF2 using the following criteria: pLDDT > 85, iPTM > 0.80, and minimal deviation of the binder backbone before versus after sequence design (RMSD < 1.5 Å). To enrich for promising backbones with reproducible sequence solutions, we prioritized backbones that yielded at least two AF2-passing sequences in this first-pass screen. Using this criterion, we selected 23 OX40L backbones (15 from Method 1 and 8 from Method 2) and 18 4-1BBL backbones (6 from each method). For each selected backbone, we then generated 100 additional sequences using ProteinMPNN to expand sequence diversity. Designs were subjected to a second, more stringent AF2 screen (pLDDT > 89, iPTM > 0.88, RMSD < 1.5 Å), yielding 26 of 2,300 OX40L designs and 97 of 1,800 4-1BBL designs that met all criteria. Finally, considering both backbone diversity and representation across design methods, we selected 18 OX40L candidates and 17 4-1BBL candidates for experimental screening. The sequences of the selected candidates and key AF2 metrics, are summarized in Supplementary Data, Supplementary Table 5, Supplementary Table6.

### Screening and characterization of masked OX40L and 41BBL

Newly designed masking domains were fused to human OX40L or human 4-1BBL using the same TEV-cleavable linker strategy used for TNFα. For OX40L, initial screening by ELISA using Expi293F culture supernatants identified 16 of 18 constructs with reduced OX40 binding before activation, and 9 of these showed increased binding after TEV cleavage (Supplementary Fig. 2a). We selected four candidates for further study, two from Method 1 (mOX40L1 and mOX40L7) and two from Method 2 (mOX40L4 and mOX40L8). AlphaFold2 models suggested that these masking domains adopt three- or four-helix-bundle folds and engage the native OX40L trimerization interface (Supplementary Fig. 2b).

After IMAC and SEC purification, all four constructs showed weak OX40 binding before activation and substantially stronger binding after TEV cleavage (Fig. 2b, Supplementary Fig. 2c). Method 2 designs showed larger EC50 shifts than Method 1 designs (Supplementary Table 7). Analytical SEC further indicated TEV-dependent reassembly of higher-order OX40L species for mOX40L4 and mOX40L8, consistent with homotrimer formation (Fig. 2c).

For 4-1BBL, ELISA-based receptor binding was not useful for the initial screen because most designs showed similar binding before and after TEV treatment (data not shown), suggesting that the monomer retains sufficient receptor-binding surface for detectable binding without trimer assembly^26^. We therefore used analytical SEC as the primary screen and prioritized designs with substantial monomer populations. All designs showed a detectable monomer peak, with estimated monomer fractions ranging from 49% to 95% (Supplementary Fig. 2d). Based on monomer content and representation across design strategies, we selected five candidates for further study: one from Method 1 (m4-1BBL2), three from Method 2 (m4-1BBL5, m4-1BBL6 and m4-1BBL8), and one from Method 3 (m4-1BBL13). AF2 models suggested that m4-1BBL2 adopts a four-helix-bundle fold, whereas the others adopt three-helix bundles, and that all engage the native 4-1BBL trimerization interface (Supplementary Fig. 2e).

After purification, all five constructs again showed similar 4-1BB binding before and after TEV treatment (Supplementary Fig. 2f). In contrast, analytical SEC showed TEV-dependent formation of higher-order species for all five constructs, consistent with homotrimer reassembly (Fig. 2d, Supplementary Fig. 2g). Among them, m4-1BBL6 showed the largest increase in the trimer peak, whereas the others showed more limited trimer formation despite largely complete linker cleavage (Supplementary Fig. 2h), suggesting that the masking domains can remain tightly associated with monomeric 4-1BBL after cleavage.

To test whether signaling was effectively suppressed in the monomeric state, we compared SEC-isolated monomeric and trimeric m4-1BBL6 using Jurkat-Lucia h4-1BB reporter cells. The trimeric species, generated by TEV cleavage and re-isolated by SEC, induced robust reporter activation comparable to TEV-treated 4-1BBL control. In contrast, the monomeric species failed to induce signaling even at a 30-fold higher concentration (9 µM versus 300 nM) (Fig. 2e). These results indicate that receptor binding alone is insufficient for productive signaling in this format and that trimer assembly and/or higher-order receptor clustering is required.

Scaffold-guided designs (Method 2) consistently produced three-helix-bundle masking domains related to the TNFα-derived templates, while still allowing substantial geometric variation (Fig. 2f). The superposition-based Method 3, although explored as a trial strategy, also yielded a well-matched and strongly masking design for 4-1BBL. Together, these results demonstrate that the masked TNFL platform can be extended across multiple TNFL targets through complementary AI-guided design routes.

### Application1: Antibody fusion for delivery of soluble TNFLs

Antibody fusions are an attractive strategy to improve the therapeutic index of TNFLs by combining targeted delivery with the favorable pharmacokinetics of IgG molecules. To evaluate whether our masking platform is compatible with antibody-based delivery, we used the monomer-stabilized masked TNFα design mTNFα7 to generate three trastuzumab fusion formats (Fig. 3a): a heavy-chain fusion (HC fusion), a light-chain fusion (LC fusion), and a dual fusion on both chains (HCLC fusion). In each construct, the masking domain was fused to the C-terminus of the trastuzumab heavy or light chain, followed by a TEV-cleavable linker as a model activation switch and the TNFα module. Upon TEV cleavage, TNFα is expected to dissociate from the antibody-tethered masking domain and reassemble into a trimer. Amino acid sequences are provided in Supplementary Data. We characterized these constructs by measuring TNFR2 and HER2 binding, assessing TEV-dependent TNFα reassembly, and testing biological activity.

**Figure 3.**
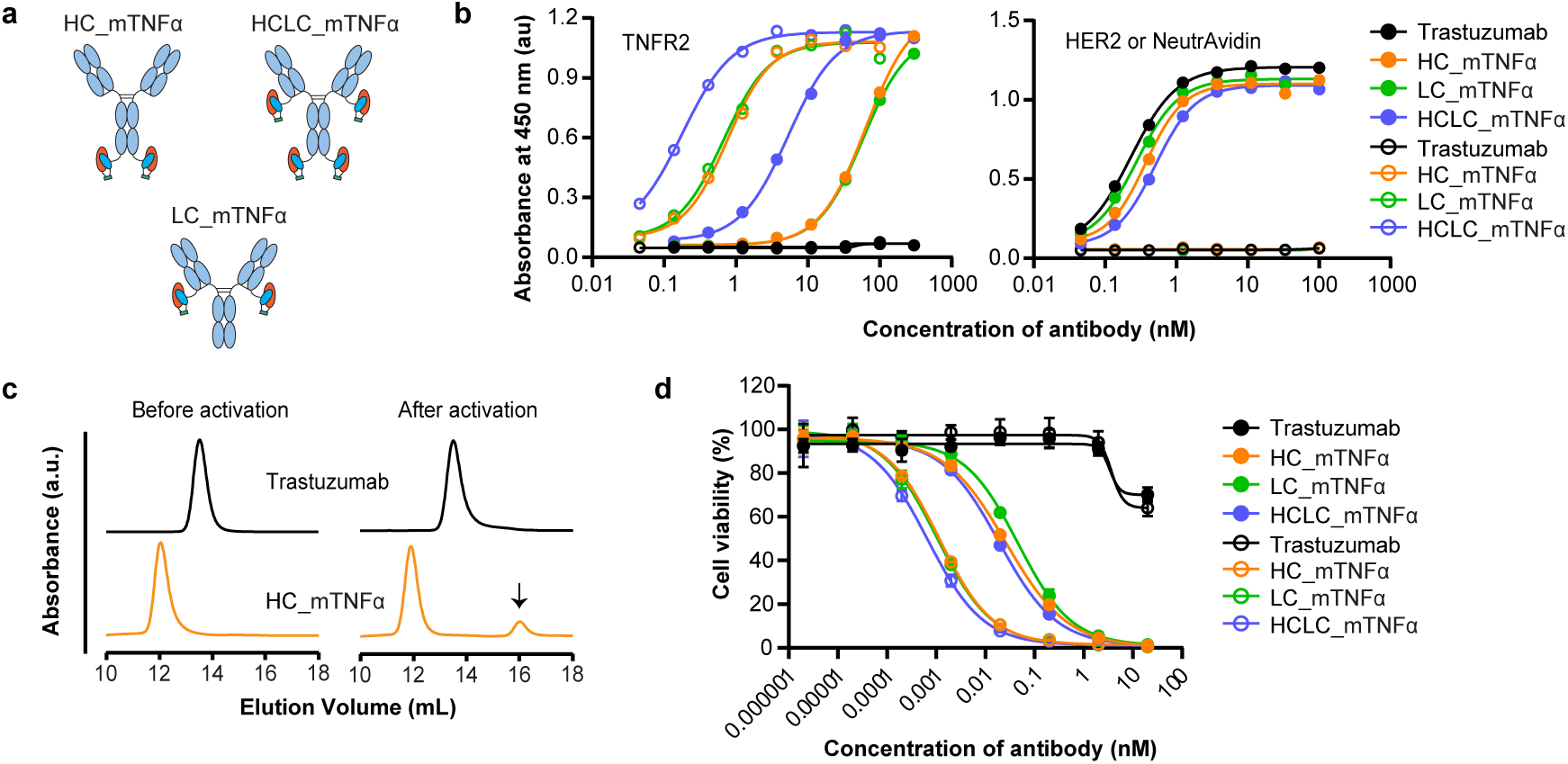
Antibody fusion enables conditional TNFα delivery in an IgG format. Masked TNFα was genetically fused to the heavy and/or light chain of the anti-HER2 antibody trastuzumab to generate IgG-format delivery molecules that retain HER2 binding while enabling protease-triggered recovery of TNFα activity. Unless otherwise noted, protein concentrations are reported per antibody molecule. a, Schematic of antibody–mTNFα fusion constructs. Masked TNFα was fused to the heavy chain (HC_mTNFα), the light chain (LC_mTNFα), or both chains (HCLC_mTNFα) of trastuzumab. In each construct, the masking domain (blue) was genetically fused to the antibody. b, ELISA binding profiles. Left, TNFR2 binding of antibody fusion proteins measured before (filled circles) and after (open circles) TEV activation. Right, HER2 binding (filled circles) and NeutrAvidin control binding (open circles) measured before activation. Data are from a single experiment and are shown as representative results. c, Oligomeric states of trastuzumab and HC_mTNFα assessed by analytical SEC before (left) and after (right) TEV activation. The arrow indicates the elution volume of the trimeric TNFα control. d, Cytotoxic activity of antibody fusion proteins in WEHI-13VAR cells before (filled circles) and after (open circles) TEV activation. Cell viability (%) was calculated as 100 × (S − B)/(V − B), where S is the sample signal, V is the 100% viability control (cells plus medium), and B is the 0% viability control (cells plus 1% Triton X-100). Data are shown as mean ± s.d. from n = 3 technical replicates.

All fusion proteins were produced in Expi293F cells and purified by Protein A affinity chromatography followed by SEC. The major SEC peak was used for downstream analyses (Supplementary Fig. 3a), and SDS–PAGE banding patterns were consistent with correct assembly of heavy and light chains for each format (Supplementary Fig. 3b).

Consistent with the masked TNFα constructs, all antibody fusions showed an activation-dependent increase in binding to human TNFR2, whereas trastuzumab alone showed no TNFR2 binding (Fig. 3b). HC and LC fusions displayed larger activation-dependent EC50 shifts (>80-fold) than the HCLC fusion (Supplementary Table 8), which may reflect avidity effects from masked TNFα in the pre-activated state. As expected, all constructs bound HER2 with similar apparent affinities, indicating that fusion of masked TNFα did not measurably impair antigen recognition (Fig. 3b, Supplementary Table 9).

Analytical SEC provided evidence for activation-dependent formation of a trimer-like TNFα species. Upon TEV treatment of the HC fusion, an additional minor peak appeared at ∼16 mL (Fig. 3c), co-eluting with the trimeric TNFα control characterized above, whereas no corresponding peak was observed for trastuzumab under the same conditions. In a WEHI-13VAR cytotoxicity assay, all antibody fusion formats showed robust masking before activation and increased activity after TEV cleavage, yielding comparable functional windows (EC50 ratio >20) across the three architectures (Fig. 3d, Supplementary Table 4). Collectively, these results show that masked TNFLs can be stably incorporated into IgG formats while preserving antigen binding, enabling protease-triggered recovery of TNFα activity and supporting a delivery mode in which functional soluble TNFα is locally released from an antibody fusion.

### Application 2: Antibody fusion for delivery of membrane-type TNFLs

Unlike TNFα, most TNF-family ligands (TNFLs), including 4-1BBL and OX40L, function primarily as membrane-associated proteins and induce signaling by crosslinking and clustering their cognate TNF receptors (TNFRs)^8^. To evaluate antibody-based delivery of such membrane-type TNFLs, we designed an asymmetric antibody format in which a single Fab arm is paired with a masked TNFL module fused to one Fc chain. In the pre-activated state, this format is expected to show limited activity because the TNFL module is masked and antigen engagement is reduced. Upon activation, TNFL-driven homotrimerization should increase functional valency, stabilize target engagement, and promote TNFR clustering and signaling (Fig. 4a).

**Figure 4.**
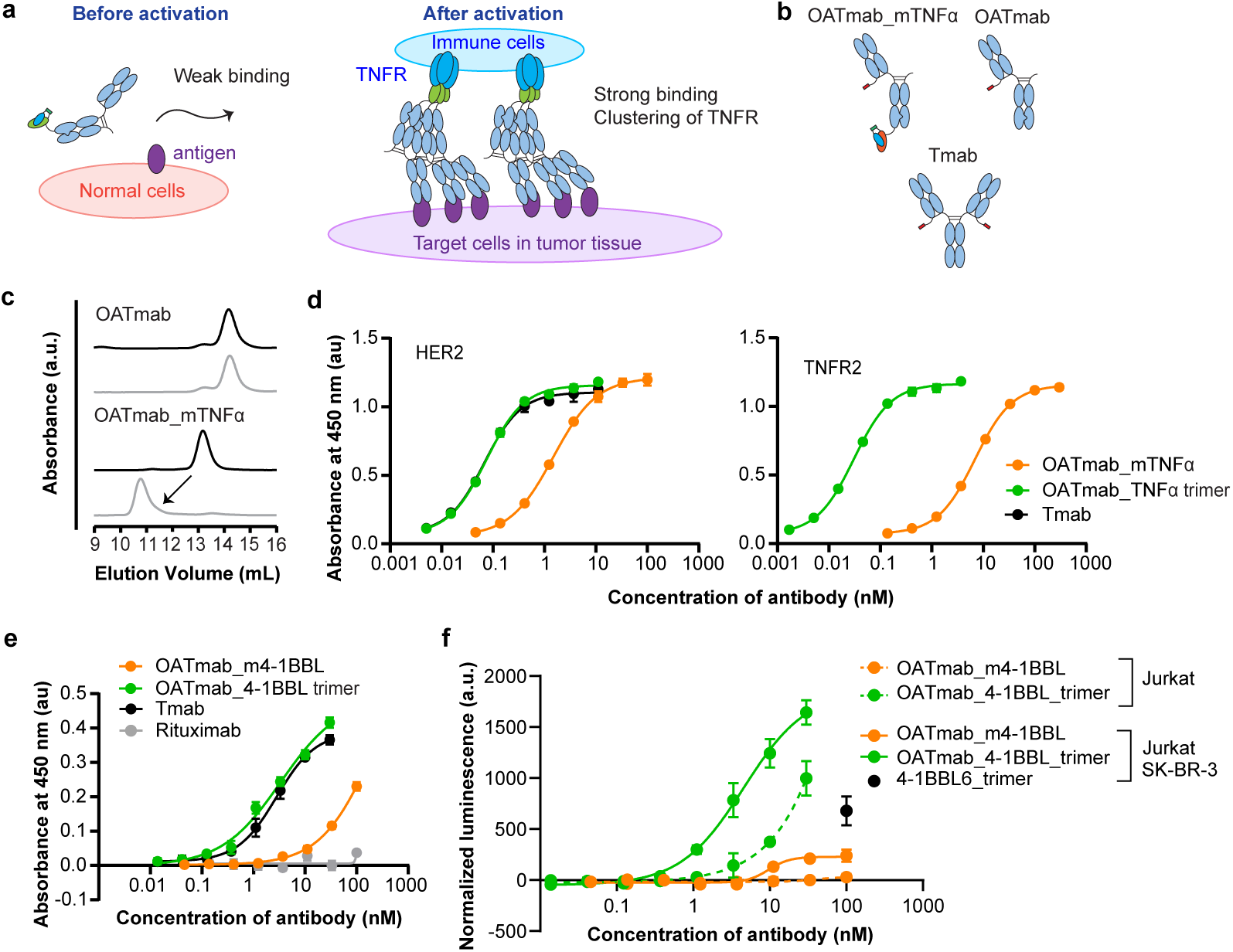
Conditional antibody multimerization for delivery of membrane-type TNFLs. A single masked TNFα or 4-1BBL module was fused to a one-armed, affinity-attenuated trastuzumab variant to generate an antibody format that undergoes activation-dependent homotrimerization. Homotrimerization was first validated using the masked TNFα construct (b–d), and functional consequences were then evaluated using an analogous masked 4-1BBL fusion in assays with HER2-positive SK-BR-3 cells and Jurkat-Lucia h4-1BB reporter cells (e, f). Protein concentrations are reported per molecular species (monomeric or trimeric antibody, as indicated). Where indicated, trimeric species were re-isolated by SEC before downstream assays. a, Proposed mechanism for delivery of membrane-type TNFLs. Before activation, the construct is predominantly monomeric with reduced apparent target engagement and masked TNFL function. Protease-triggered activation promotes assembly into a trimeric state with increased functional valency and the capacity to drive receptor crosslinking, thereby enhancing signaling. b, Design of antibody formats used to test the concept: OATmab_mTNFα, OATmab, and Tmab. All constructs are based on an affinity-attenuated trastuzumab variant^27^ and contain a C-terminal peptide tag on the light chain for site-selective drug conjugation^33^. In OATmab_mTNFα, TNFα (orange) within the masked TNFα module was fused to the C-terminus of one heavy chain. One-armed formats were assembled using knob-into-hole Fc heterodimerization mutations. c, Oligomeric states of OATmab and OATmab_mTNFα assessed by analytical SEC before (black) and after (gray) TEV activation. The arrow indicates the TEV-dependent shift in the main peak consistent with homotrimer formation. d, Binding of monomeric and trimeric antibody species to HER2 (left) and TNFR2 (right) measured by ELISA. Data are shown as mean ± s.d. from n = 3 technical replicates. e, Binding of monomeric and trimeric antibody species to HER2-positive SK-BR-3 cells measured by cell-based ELISA. Data are shown as mean ± s.d. from n = 3 technical replicates. f, Functional activity of monomeric and trimeric antibody species measured in Jurkat-Lucia h4-1BB reporter cells cultured alone (dashed lines) or co-cultured with SK-BR-3 cells (solid lines). Data are shown as mean ± s.d. from n = 3 technical replicates.

We first tested TNFL-driven antibody homotrimerization using three trastuzumab-based formats carrying previously reported HER2 affinity-attenuating mutations^27^ (Fig. 4b): a one-armed antibody fused to mTNFα7 (OATmab_mTNFα), a one-armed antibody lacking the mTNFα module (OATmab), and a conventional bivalent antibody (Tmab). In this Application 2 format, TNFα was fused directly to the Fc C-terminus to drive activation-dependent homotrimerization, whereas in Application 1 the masking domain was positioned between the antibody and TNFα (Fig. 3). TEV cleavage served as a model activation switch, and amino acid sequences are provided in Supplementary Data. All constructs were expressed in Expi293F cells and purified by Protein A chromatography followed by SEC. The major SEC peak was used for subsequent analyses (Supplementary Fig. 4a), and SDS–PAGE banding patterns were consistent with correct heavy- and light-chain assembly (Supplementary Fig. 4b).

We first assessed the oligomeric state of the antibody variants by SEC. OATmab eluted at a similar retention volume (∼14.2 mL) with or without TEV treatment, whereas OATmab_mTNFα exhibited a pronounced TEV-dependent shift, eluting ∼2.3 mL earlier after activation (Fig. 4c). Mass photometry independently confirmed an approximately threefold increase in molecular mass upon TEV treatment (Supplementary Fig. 4c), consistent with activation-dependent antibody homotrimer formation driven by TNFα assembly.

We next isolated the trimeric OATmab_TNFα species by SEC and evaluated binding to HER2 and TNFR2 by ELISA (Fig. 4d). The activated trimer showed ∼20-fold stronger apparent binding to HER2 than the pre-activated monomeric state and approached the apparent binding strength of Tmab (Supplementary Table 10). Similarly, the activated trimer showed ∼200-fold stronger apparent binding to TNFR2 than the pre-activated state (Supplementary Table 11). Together, these results demonstrate TNFL-driven homotrimerization of the antibody format and conditional enhancement of both antigen engagement and TNFR binding upon activation.

We next replaced mTNFα7 with masked 4-1BBL (m4-1BBL6) to evaluate delivery of a membrane-type TNFL. As in the TNFα case, the fusion protein was expressed in Expi293F cells and purified by Protein A chromatography followed by SEC. Following TEV activation, the homotrimeric species was re-isolated by SEC. The major SEC peak showed a TEV-dependent shift of ∼2.4 mL, consistent with trimerization, and the corresponding peak fraction was used for subsequent analyses (Supplementary Fig. 4d).

In binding assays using HER2-expressing SK-BR-3 cells, the trimeric fusion displayed stronger apparent binding than the monomeric form and achieved binding comparable to the IgG-format control (Tmab) (Fig. 4e). In Jurkat-Lucia h4-1BB reporter assays, the trimeric fusion induced stronger signaling than the monomer; notably, the strongest activation was observed when reporter cells were co-cultured with SK-BR-3 cells, consistent with localized clustering of 4-1BB driven by cell-surface presentation of the trimeric fusion on HER2-positive target cells (Fig. 4f). This interpretation was supported by competitive inhibition with an anti-HER2 antibody (Supplementary Fig. 4e).

Collectively, these data indicate that simultaneous engagement of the tumor antigen and TNFR, which is enabled by conditional antibody homotrimerization, is critical for efficient signaling by membrane-type TNFL delivery formats.

### Application 3: An inducible multimerizing ADC format with an expanded therapeutic window

Antibody–drug conjugates (ADCs) are an established class of targeted anti-cancer therapeutics that can achieve high potency while limiting systemic toxicity^28, 29^. Efficient internalization is a key determinant of ADC efficacy, and receptor crosslinking and avidity have been reported to enhance internalization for multiple receptor targets^30, 31, 32^. We hypothesized that the switchable antibody homotrimerization mediated by masked TNF ligands could be repurposed to create an ADC architecture in which protease-triggered assembly increases functional valency and payload density in a target-dependent manner, thereby expanding the therapeutic window. The proposed mechanism is summarized in Fig. 5a: prior to activation, the construct is predominantly monomeric with reduced apparent avidity and low drug-to-antibody ratio (DAR), whereas activation induces trimerization, increasing effective valency, apparent binding strength, and payload density.

**Figure 5.**
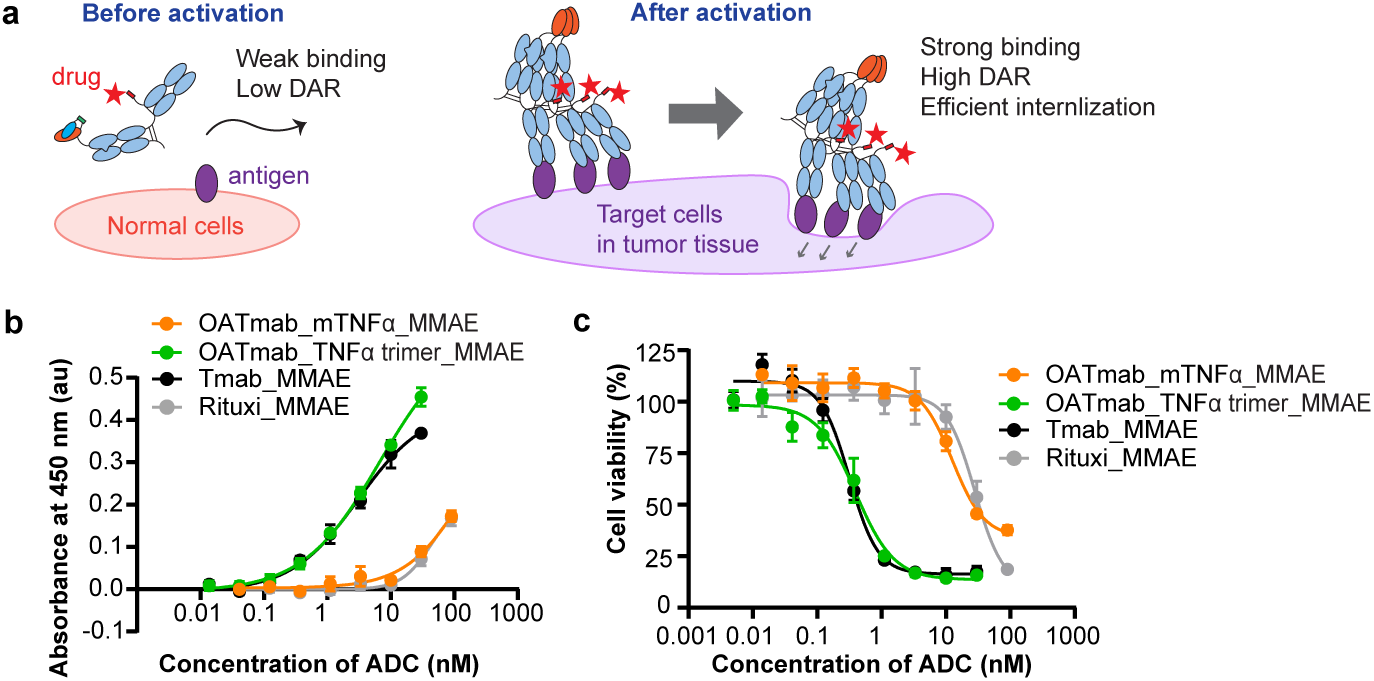
Masked TNFα enables a conditionally multimerizing ADC concept with an expanded therapeutic window. The antibody formats shown in Fig. 4b were conjugated with MMAE, and cell binding and cytotoxicity were evaluated in HER2-positive SK-BR-3 cells. Protein concentrations are reported per molecular species (monomeric or trimeric ADC, as indicated). Where indicated, trimeric species were re-isolated by SEC before downstream assays. a, Proposed mechanism for the switchable ADC concept. Before activation, the construct is predominantly monomeric with reduced apparent binding and low payload density. Protease-triggered activation promotes assembly into a trimeric state with increased functional valency and complex-level payload density, which may enhance target-cell killing. b, Binding of monomeric and trimeric ADC species to HER2-positive SK-BR-3 cells measured by cell-based ELISA. Data are shown as mean ± s.d. from n = 3 technical replicates. c, Cytotoxic activity of ADCs in HER2-positive SK-BR-3 cells. Cell viability (%) was calculated as 100 × S/V, where S is the sample signal and V is the 100% viability control (cells plus medium). Data are shown as mean ± s.d. from n = 3 technical replicates.

To test this concept, we used the antibody formats shown in Fig. 4b, which include a C-terminal light-chain conjugation tag (GSGSGSAPPLPPRNRPRLLEVLFQGPSGSGKGSGSYAAA) enabling site-selective conjugation of endo-BCN–Val–Cit–PAB–MMAE^33^. The drug linker was covalently conjugated to the antibody formats, and a trimeric ADC was generated by TEV activation of the monomeric ADC followed by SEC purification. Homotrimerization of ADC was confirmed by SEC, consistent with the behavior of the corresponding unconjugated antibodies (Supplementary Fig. 5a). Intact-MS analysis estimated the drug-to-antibody ratios (DARs) of OATmab_mTNFα_MMAE, OATmab_TNFα trimer_MMAE, Tmab_MMAE, and Rituximab_MMAE to be 0.34, 0.60, ∼2.0 and ∼2.0, respectively.

In a cell-based binding assay, the trimeric ADC showed stronger apparent binding than the monomeric ADC and approached the apparent binding of the bivalent ADC (Fig. 5b).

Consistently, in cytotoxicity assays the trimeric ADC exhibited ∼30-fold greater potency than the monomeric ADC (Fig. 5c, Supplementary Table 12). The corresponding unconjugated trimeric antibody showed no detectable cytotoxicity (Supplementary Fig. 5b). Moreover, cytotoxicity of the trimeric ADC was competitively inhibited by excess unconjugated antibody (Tmab), indicating that killing was HER2-dependent (Supplementary Fig. 5c). Notably, the trimeric ADC achieved cytotoxic efficacy comparable to the bivalent ADC despite a modestly lower DAR, suggesting that activation-dependent multimerization can compensate for reduced payload loading through increased apparent avidity and/or target engagement, which potentially leaded to enhanced internalization.

Together, these results demonstrate that masked TNFα can drive activation-dependent homotrimerization of an ADC, thereby enhancing apparent binding and target-cell killing.

## Discussion

In this study, we used AI-enabled protein design to create de novo masking domains that stabilize TNF-family ligands (TNFLs) in an inactive monomeric state by targeting their native trimerization interfaces. Proteolytic activation released the masking domains, promoted homotrimer reassembly, and restored biological activity. This strategy was generalizable across TNFα, OX40L, and 4-1BBL, and the resulting masked TNFLs were compatible with both full-length IgG and one-armed antibody formats while preserving antigen binding. Beyond TNFL delivery, this switchable self-assembly mechanism also enabled a conditionally multimerizing ADC architecture with the potential to improve therapeutic index.

A central challenge in TNFL therapeutics is to achieve localized activation while retaining compatibility with IgG/Fc architectures that enable targeted delivery and provide favorable pharmacokinetics and manufacturability. Previous approaches have only partially addressed these requirements. Receptor-based masking strategies can provide conditional activation^17, 18^, but obligate trimeric architectures are difficult to integrate into conventional IgG/Fc formats. Conversely, attenuated-cytokine delivery strategies can improve selectivity but may alter ligand biology^19^. Split-ligand antibody formats or single chain TNFLs improve IgG compatibility^34, 35^, yet constitutive TNFL activity may still increase the risk of on-target effects in normal tissues. In contrast, our masked TNFLs combine protease-triggered activation with stabilization of an inactive monomeric state that is intrinsically compatible with IgG/Fc architectures, offering a distinct and potentially safer strategy to couple conditional activation with antibody-based TNFL delivery.

Our results also highlight the value of AI-enabled design for targeting a native trimerization interface that is experimentally difficult to access. Although interface-binding peptides and protein domains have previously been reported for TNFα^36, 37, 38^, our attempts to repurpose such binders as masking modules were unsuccessful because they did not provide the required balance of solubility, stability, and post-cleavage reversibility (Supplementary Fig. 6). By contrast, computational design enabled direct exploration of diverse backbone topologies and binding modes around a specified interface^20, 21, 22^, and supported extension to related TNFLs through scaffold-guided and superposition-based workflows. The recovery of multiple responsive designs from relatively small experimental screens underscores the practical versatility of this approach.

Beyond cytokine delivery, masked TNFLs enabled an ADC concept based on gated multivalency. In this format, protease-triggered TNFL self-assembly promotes antibody homotrimerization, increasing functional valency and complex-level payload density relative to the predominantly monomeric pre-activated state. This architecture may also enhance receptor crosslinking and subsequent internalization, in line with reports that receptor crosslinking promotes ADC uptake^30, 31, 32^. Mechanistically, this differs from conditional ADC strategies such as Probody formats^39, 40^, which primarily gate antigen recognition by masking the paratope. Our design instead introduces an activation-dependent amplification step at the level of the assembled complex. In addition, payload-mediated cytotoxicity from such ADCs could potentially be combined with cancer immunotherapies^41, 42^. Although any reduction in off-target toxicity will require experimental validation, the low-payload pre-activated state and locally assembled trimer may provide a route to improving therapeutic index in protease-rich tumors.

Important next steps will be to establish in vivo proof of concept for masked TNFLs and antibody fusion formats, including validation of tumor-selective activation using circulation-stable protease linkers together with integrated PK/PD analyses. For ADC applications, it will then be important to define the mechanistic basis of efficacy, including the contributions of activation-dependent multivalency, receptor clustering, internalization, trafficking, and payload density. More broadly, further development of this platform will require optimization of masking strength versus post-cleavage reversibility, as well as evaluation of developability and immunogenicity for de novo masking domains. Together, these results establish a general framework for controlling TNFL assembly and function, and expand the design space for conditional immunomodulatory therapeutics.

## Methods

### Computational design of masking domains for TNFα

Masking domains for human TNFα were designed using a computational pipeline consisting of RFdiffusion for backbone generation^22^, ProteinMPNN for sequence design^23^, and AlphaFold2 (AF2) for complex prediction^24^ and in silico screening (Fig. 1a).

#### Backbone generation (RFdiffusion)

The human TNFα trimer structure (PDB ID: 1A8M) was used as the starting model. A monomeric target was prepared in PyMOL by retaining chain A, deleting chains B and C, and removing the first five N-terminal residues (VRSSS). RFdiffusion was then run in protein–protein interaction design mode to generate 1,000 candidate binder backbones targeting a hydrophobic hotspot on TNFα defined by residues L52, Y54, P112, Y114, I150, and L152. Candidate backbones were filtered for compactness by requiring the binder radius of gyration (Rg) to fall below a length-dependent threshold defined as 2.6 × (binder length)^0.365^, yielding 182 backbones for sequence design.

#### Sequence design (ProteinMPNN)

For each filtered backbone, 10 sequences were generated in fixed-backbone mode using ProteinMPNN, with cysteine excluded from the design alphabet.

#### Structure prediction and screening (AlphaFold2)

AF2 was run in single-sequence mode without MSA, using TNFα as a fixed custom template, and one model was generated for each designed sequence. Designs were prioritized based on mean pLDDT (>85), iPTM (>0.85), and binder RMSD (<1.5 Å) relative to the designed backbone. Twenty-one designs were selected for experimental screening.

### Scaffold-guided design

Artificial monomer structures of OX40L and 4-1BBL were prepared from the trimer structures (PDB ID: 2HEV for OX40L and PDB ID: 6BWV for 4-1BBL) by retaining a single protomer and deleting the remaining chains in PyMOL. Hotspot residues were defined as L45, L81, V83, I114, and I116 for OX40L, and F4, Y54, F56, F111, L115, and F150 for 4-1BBL. The TNFα masking domains mTNFα7 and mTNFα8 were used as scaffolds to guide backbone generation. Secondary-structure and adjacency inputs were generated with the make_secstruc_adj.py script provided in the RFdiffusion repository. For each target ligand, scaffold-guided RFdiffusion was run using either mTNFα7 or mTNFα8 and directed toward the predefined hotspot residues. Fifty backbones were generated per scaffold, yielding 100 total backbones per target TNFL. Sequence design and AF2-based screening were performed as described above.

### Superposition-based design

For 4-1BBL, the TNFα protomer in the mTNFα7 or mTNFα8 complex was structurally aligned to the artificial 4-1BBL monomer in PyMOL. The TNFα chain was then removed, and the TNFα masking domain was merged with the 4-1BBL monomer to generate a starting complex.

RFdiffusion partial diffusion^21^ was applied to each starting complex to generate 50 backbone variants, yielding 100 total backbones for 4-1BBL. Sequence design and AF2-based screening were performed as described above.

### Expression and purification of proteins

All constructs were cloned into the pTwist cDNA 3.4 mammalian expression vector (Twist Bioscience) and transiently expressed in Expi293F cells (Thermo Fisher Scientific) using a PEI-based transfection protocol adapted from a published method^43^. Cells were transfected at a density of 2.5 × 10^6^ cells/mL with PEI MAX (Kyfora Bio) and plasmid DNA. At 16–20 h post-transfection, sodium propionate, valproic acid, and glucose were added to enhance protein expression. Cultures were harvested 48–72 h later by centrifugation (3,000 × g, 15 min), and clarified supernatants were used for screening and purification.

His-tagged TNFL and masked TNFL constructs were purified using PROTEINDEX Ni-Penta Agarose 6 Fast Flow resin (Marvelgent Biosciences). The resin was washed with PBS supplemented with 300 mM NaCl, and bound proteins were eluted with PBS containing 300 mM NaCl and 250 mM imidazole. Antibody and antibody-fusion proteins were purified using Protein A agarose resin (GoldBio), washed with PBS, and eluted with 0.1 M sodium acetate (pH 3.5), followed by immediate neutralization with 1 M Tris–HCl (pH 9.0).

All proteins were further purified by size-exclusion chromatography (SEC). Samples were concentrated using Amicon Ultra centrifugal filters (MilliporeSigma) and applied to a Superdex 200 Increase 10/300 GL column (Cytiva) equilibrated in SEC buffer (20 mM HEPES, 150 mM NaCl, 5% (w/v) sorbitol, pH 7.4). Peak fractions were collected for subsequent characterization. Protein concentrations were determined from absorbance at 280 nm using sequence-derived molar extinction coefficients^44^.

### Activation of masked TNFLs by TEV protease

Masked TNFL constructs were activated by cleavage of the TEV-sensitive linker using either in-house–prepared TEV protease or commercial cold-active TEV protease (NEB). Unless otherwise noted, reactions were performed in TEV cleavage buffer (50 mM Tris–HCl, 150 mM NaCl, 0.5 mM EDTA, pH 8.0) under conditions empirically determined to achieve near-complete linker cleavage. Representative conditions were 30 °C for 1 h with TEV protease for ELISA and analytical SEC, or cold-active TEV protease for cell-based assays. Activated samples were used directly in downstream assays unless SEC re-isolation of the homotrimeric species is explicitly stated.

Linker cleavage efficiency was assessed by SDS–PAGE using Mini-PROTEAN TGX Stain-Free precast gels (Bio-Rad) and was considered near-complete when the higher-molecular-weight fusion band was largely converted to the expected lower-molecular-weight cleavage products with minimal residual uncleaved species.

### Enzyme Linked Immunosorbent Assay (ELISA)

High-binding 96-well plates (Nunc MaxiSorp; Thermo Fisher Scientific) were coated overnight at 4 °C with NeutrAvidin (1 µg/mL in PBS, 50 µL/well), washed with PBS containing 0.05% Tween-20 (PBS-T), and blocked with Blocker Casein in PBS (Thermo Fisher Scientific) for 1 h at room temperature. Biotinylated target proteins (TNFR2, OX40, 4-1BB, and HER2; ACROBiosystems) were captured at 0.5 µg/mL in PBS (50 µL/well) for 30 min at room temperature. After washing, protein samples (± TEV activation, as described above) were added (50 µL/well) and incubated for 30 min at room temperature. Bound proteins were detected with either Direct-Blot HRP anti-DYKDDDDK (FLAG) antibody (BioLegend) for FLAG-tagged constructs or Peroxidase AffiniPure goat anti-human IgG (Fcγ fragment-specific; Jackson ImmunoResearch) for antibody formats, followed by development with TMB substrate (BioLegend) and quenching with 1 N HCl. Absorbance at 450 nm was measured using a SpectraMax M3 plate reader (Molecular Devices). Dose–response curves and EC50 values were determined in GraphPad Prism v10.6.1 using a four-parameter logistic model.

For the Adalimumab-binding assay, plates were coated directly with Adalimumab (1 µg/mL in PBS) instead of using NeutrAvidin-mediated capture. TNFα or masked TNFα samples (± TEV activation) were then added and detected with HRP anti-FLAG antibody as described above.

### Analytical Size Exclusion Chromatography (SEC)

Molecular size and oligomeric state were assessed by analytical size-exclusion chromatography using a Superdex 200 Increase 10/300 GL column (Cytiva) on an ÄKTA FPLC system. The column was equilibrated with analytical SEC buffer (20 mM HEPES, 150 mM NaCl, pH 7.4) at 4 °C. Protein samples (250 µL at 5 or 10 µM, as indicated) were injected and eluted at a flow rate of 0.5 mL/min, and absorbance at 280 nm was monitored. Chromatograms are shown in arbitrary units.

### Cell killing activity of masked TNFα

The biological activity of masked TNFα was evaluated using WEHI-13VAR murine fibrosarcoma cells (ATCC). Cells were maintained in RPMI-1640 supplemented with 10% (v/v) fetal bovine serum (FBS) at 37 °C in a humidified incubator with 5% CO₂. For assays, cells were detached with TrypLE Express Enzyme (Thermo Fisher Scientific), resuspended in RPMI-1640 containing 10% FBS, and adjusted to 2.0 × 10⁵ cells/mL. Cell suspensions (50 µL; 10,000 cells per well) were plated in 96-well clear-bottom plates (CELLSTAR; Greiner) and pre-incubated for 6 h at 37 °C.

Masked TNFα samples (± TEV activation, as described above) were diluted in RPMI-1640 containing 10% FBS and 1 µg/mL actinomycin D, and 10-fold serial dilutions were prepared in the same medium. After pre-incubation, 50 µL of each dilution was added to the cells (final volume, 100 µL per well), yielding a maximum final TNFα concentration of 20 nM and a final actinomycin D concentration of 500 ng/mL. Vehicle controls received medium containing 10% FBS and 1 µg/mL actinomycin D, and 100% killing controls received the same medium supplemented with 2% (v/v) Triton X-100. Plates were incubated for a further 16–18 h at 37 °C.

Cell viability was measured using CellTiter-Glo 2.0 (Promega) according to the manufacturer’s instructions. Luminescence was recorded using a SpectraMax M3 plate reader (Molecular Devices). Relative viability was calculated from luminescence values as described in the figure legends. Dose–response curves and EC50 values were determined in GraphPad Prism v10.6.1 using a four-parameter logistic model.

### Cell functional activity of masked 4-1BBL

The biological activity of masked 4-1BBL was evaluated using Jurkat-Lucia h4-1BB reporter cells (InvivoGen). Cells were maintained and assayed according to the manufacturer’s instructions. Briefly, cells were cultured in IMDM supplemented with 10% (v/v) fetal bovine serum (FBS), 100 U/mL penicillin–streptomycin, and Normocin at 37 °C in a humidified incubator with 5% CO₂. Selection antibiotics (blasticidin and zeocin) were added at the recommended concentrations every other passage.

For assays, cells were resuspended in assay medium (IMDM containing 10% FBS and 100 U/mL penicillin–streptomycin) at 5.6 × 10⁵ cells/mL and dispensed into 96-well plates (180 µL per well; CELLSTAR, Greiner). Samples were prepared as 3-fold serial dilutions in PBS and added at 20 µL per well. The maximum final concentrations of monomeric and trimeric m4-1BBL6 were 9,000 nM and 300 nM, respectively. TEV-treated 4-1BBL was included as a control at a final concentration of 100 nM. Plates were incubated for 24 h at 37 °C.

Reporter activity was quantified by transferring 20 µL of culture supernatant to white opaque 96-well plates (Thermo Fisher Scientific), adding 50 µL of QUANTI-Luc-4 reagent (InvivoGen), and measuring luminescence on a SpectraMax M3 plate reader (Molecular Devices). Data were analyzed in GraphPad Prism v10.6.1.

### Mass photometry

Mass photometry was used to assess the oligomeric state of OATmab_mTNFα before and after TEV activation. Measurements were performed on a Refeyn mass photometer according to the manufacturer’s instructions. The instrument was calibrated on the day of measurement using protein standards of known molecular mass (Thermo Fisher Scientific) prepared in the same buffer as the samples.

Immediately before measurement, samples were diluted to ∼15 nM in mass photometry buffer (50 mM Tris–HCl, 150 mM NaCl, 0.5 mM EDTA, pH 8.0). Movies were recorded for 60 s at room temperature. Single-molecule landing events were analyzed using Refeyn Acquire and Discover software with default settings, and particle contrast values were converted to molecular mass using the calibration curve. Events with a fit error <0.35 were retained for analysis. Mass distributions were plotted as histograms with a 5 kDa bin width, and peak masses were determined by Gaussian fitting in GraphPad Prism v10.6.1.

### Cell functional activity of antibody-fused masked 4-1BBL

The biological activity of antibody-fused masked 4-1BBL was evaluated using Jurkat-Lucia h4-1BB reporter cells in the presence or absence of HER2-positive SK-BR-3 cells (ATCC). SK-BR-3 cells were maintained in McCoy’s 5A medium supplemented with 10% (v/v) fetal bovine serum (FBS) at 37 °C in a humidified incubator with 5% CO₂. For assays, SK-BR-3 cells were detached with TrypLE Express (Thermo Fisher Scientific), resuspended in growth medium containing 10% FBS, and seeded at 2.0 × 10⁴ cells per well in 96-well clear-bottom plates (CELLSTAR; Greiner). For conditions without SK-BR-3 cells, growth medium alone was added. Plates were incubated for 24 h at 37 °C.

The next day, 20 µL of protein samples prepared in PBS was added to each well (final concentrations: 100 nM monomeric species or 30 nM trimeric species, as indicated) and incubated for 30 min at 37 °C. Jurkat-Lucia h4-1BB cells were resuspended in assay medium (IMDM containing 10% FBS and 100 U/mL penicillin–streptomycin) at 7.7 × 10⁵ cells/mL, and 130 µL was added to each well. Plates were incubated for a further 24 h at 37 °C.

Reporter activity was quantified from 20 µL of culture supernatant as described above. Dose–response curves and EC50 values were determined in GraphPad Prism v10.6.1 using a four-parameter logistic model.

### Production of antibody-drug conjugates (ADCs)

endo-BCN-Val-Cit-PAB-MMAE (BroadPharm) was enzymatically conjugated to tyrosine residues within the C-terminal tag2 sequence on the antibody light chain (Supplementary Data), following a previously described procedure^33^. Antibody (5 µM), endo-BCN-Val-Cit-PAB-MMAE (50 µM), and tyrosinase (D42; 250 nM) were mixed in reaction buffer (50 mM phosphate, pH 6.5, 2 µM CuSO₄, 5% DMSO) and incubated at 25 °C for 4 h with shaking at 500 rpm. Reactions were quenched by addition of tropolone to a final concentration of 4 mM.

Reaction mixtures were buffer-exchanged into PBS by SEC (Superdex 200 Increase 10/300 GL). To generate trimeric ADCs, monomeric ADCs were incubated with cold-active TEV protease at 4 °C overnight, followed by SEC to isolate the trimeric species. Samples were then exchanged into 10 mM acetate buffer (pH 5.5) containing 5% (w/v) sorbitol for cell-based assays, or into 50 mM ammonium acetate (pH 6.5) for intact mass spectrometry, using Amicon Ultra centrifugal filters (MilliporeSigma).

### Intact mass spectrometry and DAR estimation

Intact mass spectrometry was performed at the Metabolomics and Proteomics (MAP) Core at the University of North Carolina. Samples were reduced with 10 mM DTT at 37 °C for 30 min and diluted to 0.5 µg/µL in peptide background electrolyte. Samples were analyzed using a ZipChip system (HR Chip; 908 Devices) interfaced to a Q Exactive HF Biopharma mass spectrometer (Thermo Fisher Scientific). Spectra were acquired in positive-ion mode over m/z 1000–4000 with in-source CID (50 eV) and deconvoluted using UniDec. The fraction of conjugated light chain species was estimated from deconvoluted peak heights as:

% conjugated LC = Iconjugated / (Iconjugated + Iunconjugated),

where I denotes the deconvoluted peak height of each light-chain species. DAR was then estimated as:

DAR = n × (% conjugated LC),

where n is the number of light chains in the construct (e.g., n = 2 for IgG formats and n = 1 for one-armed formats).

### Cell binding activity of antibodies and ADCs (cell-based ELISA)

Cell binding activity of antibodies and ADCs was evaluated using SK-BR-3 human breast cancer cells (ATCC). Cells were maintained as described above. For assays, cells were detached with TrypLE Express (Thermo Fisher Scientific), resuspended in growth medium containing 10% FBS, and seeded at 2.0 × 10^4^ cells per well in poly-L-lysine–coated 96-well clear-bottom plates (CELLSTAR; Greiner). Plates were incubated for 24 h at 37 °C.

Samples were prepared as 3-fold serial dilutions in ice-cold PBS containing 1% (w/v) BSA. After removal of culture medium, cells were washed once with ice-cold PBS, and 50 µL of each dilution was added (maximum final concentration, 30–100 nM, as indicated). Plates were incubated on ice for 45 min to allow binding while minimizing internalization. Bound antibodies were detected with Peroxidase AffiniPure Goat anti-human IgG (Fcγ fragment-specific; Jackson ImmunoResearch) diluted in ice-cold PBS containing 1% BSA, followed by TMB development and quenching with 1 N HCl. Absorbance at 450 nm was measured using a SpectraMax M3 plate reader (Molecular Devices). Data were analyzed in GraphPad Prism, and EC50 values were obtained by nonlinear regression using a four-parameter logistic model.

### Cytotoxic activity of ADCs

Cytotoxic activity of ADCs was evaluated using SK-BR-3 human breast cancer cells (ATCC). Cells were maintained as described above. For assays, cells were detached with TrypLE Express (Thermo Fisher Scientific), resuspended in growth medium containing 10% FBS, and seeded at 2,700 cells per well in 96-well clear-bottom plates (CELLSTAR; Greiner). Plates were incubated for 24 h at 37 °C.

ADC or protein samples were prepared as 3-fold serial dilutions in growth medium containing 10% FBS and added at 10 µL per well, yielding final sample concentrations of up to 30 or 90 nM, as indicated. Vehicle controls received growth medium alone. Plates were incubated for a further 3 days at 37 °C. For competition assays, trimeric ADC was tested at 1 nM in the presence of increasing concentrations of unconjugated Tmab (up to 100 nM).

Cell viability was measured using CellTiter-Glo 2.0 (Promega) as described above. Relative viability was calculated from luminescence values as described in the figure legends. Data were analyzed in GraphPad Prism, and IC50 values were obtained by nonlinear regression using a four-parameter logistic model.

## Data Availability

All amino acid sequences of the proteins evaluated in this study are provided in Supplementary Data. All other data supporting the findings of this study are available within the paper and its Supplementary Information or from the corresponding author upon reasonable request.

## Acknowledgements

We thank the Metabolomics and Proteomics (MAP) Core at the University of North Carolina at Chapel Hill for support with intact mass spectrometry analyses. We are grateful to McGuire Metts for technical guidance on protein design methods and to Christopher Shelby for technical support with ADC synthesis. S.K. acknowledges support from Daiichi Sankyo during his research visit to the University of North Carolina at Chapel Hill. ChatGPT (OpenAI) was used for language editing of the manuscript. All outputs were reviewed by the authors, who take full responsibility for the final content. Some schematics were created in BioRender and finalized in Adobe Illustrator. Created in BioRender. Kudo, S. (2026). BioRender.com/2riypzp. This work was supported by a NIH grant to B.K. (R35-GM131923).

## Author Contributions

S.K. conceived the project, designed and performed the experiments, acquired and analyzed the data, and wrote the manuscript. B.K. supervised the study and reviewed and edited the manuscript. Both authors discussed the results and approved the final manuscript.

## Competing Interests

S.K. is an employee of Daiichi Sankyo. The University of North Carolina at Chapel Hill plans to file a patent application related to this work, with S.K. and B.K. as inventors. The authors declare no other competing interests.

## Notes

### Competing Interest Statement

The authors have declared no competing interest.

