## Supplementary Information for "De novo masking domains that gate TNF-family ligand assembly and activity"

This Word file includes:

Supplementary Figures 1-6

Supplementary Tables 1-12

Additional data file:

Supplementary Data includes amino acid sequences of protein constructs in this study.

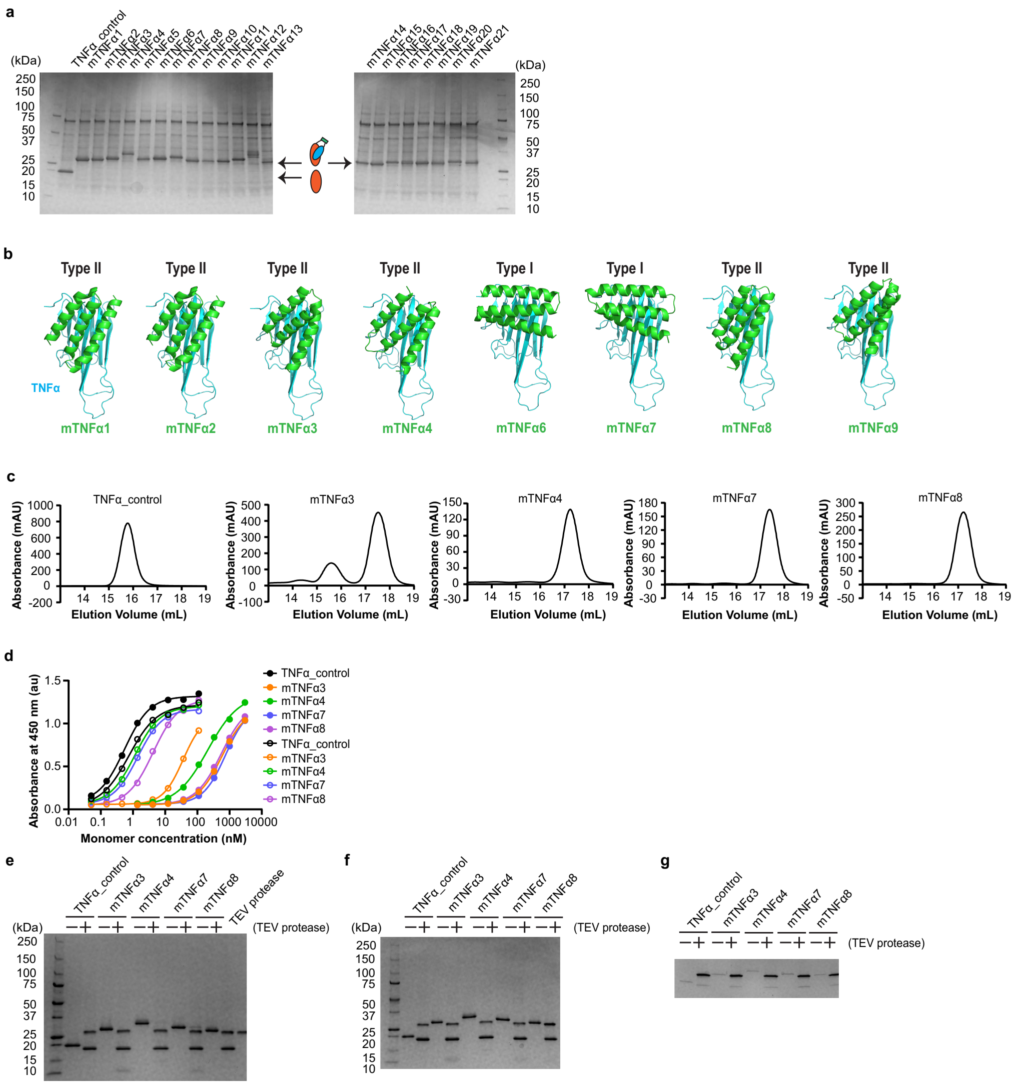

**Supplementary Figure 1. Computational design, screening and characterization of masked TNFα.**

a, SDS–PAGE of Expi293F culture supernatants expressing designed masked TNFα candidates under non-reducing conditions. Culture supernatant (10 μL) was loaded per lane. Upper and lower arrows indicate masked TNFα candidates and the TNFα_control, respectively. b, AlphaFold2-predicted complex structures of hit clones (mTNFα1, mTNFα2, mTNFα3, mTNFα4, mTNFα6, mTNFα7, mTNFα8 and mTNFα9). TNFα is shown in cyan and masking domains in green. Type I masking domains are oriented approximately perpendicular to the TNFα β-sheet surface, whereas Type II masking domains are oriented more parallel to the β-sheet surface. c,

SEC chromatograms from preparative purification of TNFα_control and selected masked TNFα constructs. Main peaks were collected for downstream analyses. Under these conditions, TNFα_control eluted earlier than all masked TNFα constructs. d, Adalimumab binding of selected masked TNFα constructs measured by ELISA before (filled circles) and after (open circles) TEV activation. Data are from a single experiment and are shown as representative results. e, SDS–PAGE of ELISA samples with and without TEV treatment under non-reducing conditions. Samples (10 µL of 3 µM) were loaded per lane. Near-complete linker cleavage was assessed by the expected band shift of the fusion protein. TEV protease bands (~27 kDa) are present in all TEV-treated samples. f, SDS–PAGE of analytical SEC samples with and without TEV treatment under non-reducing conditions. Samples (4 µL of 10 µM) were loaded per lane. Near-complete linker cleavage was confirmed by the expected band shift. TEV protease bands (~27 kDa) are present in all TEV-treated samples. g, SDS–PAGE of functional assay samples with and without TEV treatment under non-reducing conditions. Samples (10 µL of 400 nM) were loaded per lane. Near-complete linker cleavage was confirmed by the expected band shift. TEV protease bands are present in all TEV-treated samples.

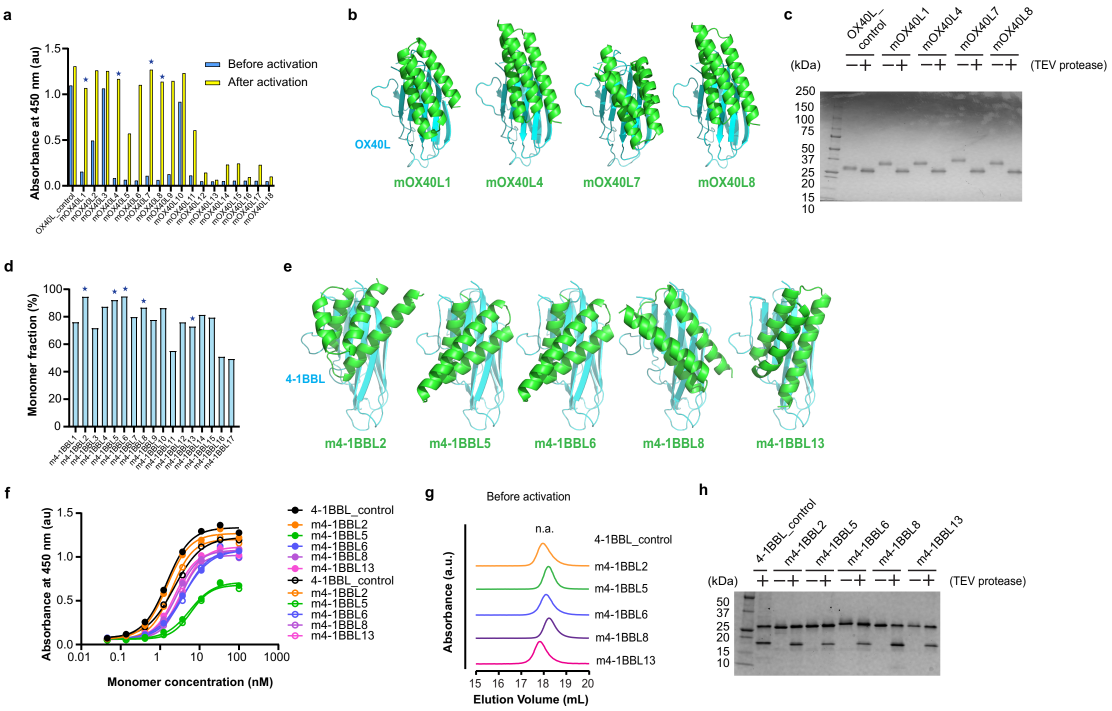

**Supplementary Figure 2. Design, screening and characterization of masked OX40L and 4-1BBL.**

a, Initial ELISA screen of masked OX40L using plate-immobilized OX40 as the target. Signals before (blue) and after (yellow) TEV activation are shown. Asterisks indicate hit clones selected for further characterization. b, AlphaFold2-predicted complex structures of representative hit clones (mOX40L1, mOX40L4, mOX40L7 and mOX40L8). OX40L is shown in cyan and the masking domain in green. c, SDS–PAGE of ELISA samples with and without TEV treatment under non-reducing conditions. Samples (10 µL of 3 µM) were loaded per lane. Near-complete linker cleavage was assessed by the expected band shift. TEV protease (~27 kDa) co-migrates with OX40L-derived bands in TEV-treated samples. d, Initial screen of masked 4-1BBL by analytical SEC. Monomer fraction (%) was estimated as 100 × M/(O + M), where M is the baseline-corrected absorbance of the monomer peak and O is the summed baseline-corrected absorbance of higher-order peaks. Asterisks indicate hit clones selected for further characterization. e, AlphaFold2-predicted complex structures of representative hit clones (m4-

1BBL2, m4-1BBL5, m4-1BBL6, m4-1BBL8 and m4-1BBL13). 4-1BBL is shown in cyan and the masking domain in green. f, 4-1BB binding of selected masked 4-1BBL constructs measured by ELISA before (filled circles) and after (open circles) TEV activation. Data are from a single experiment and are shown as representative results. g, Oligomeric state of selected masked 4-1BBL constructs assessed by analytical SEC before TEV activation. h, SDS–PAGE of SEC samples with and without TEV treatment under non-reducing conditions. Samples (4 μL of 10 μM) were loaded per lane. Near-complete linker cleavage was assessed by the expected band shift. TEV protease bands (~27 kDa) are present in all TEV-treated samples.

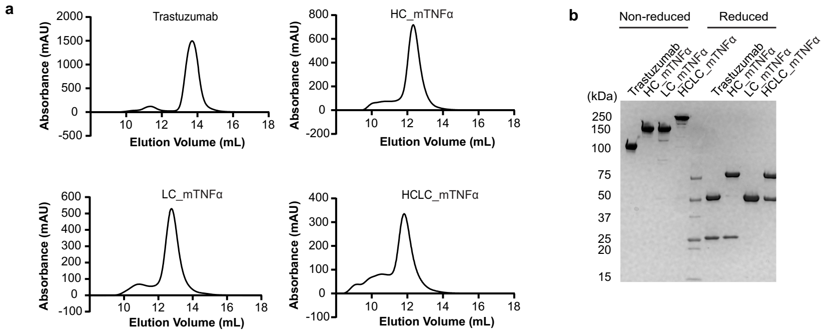


**Supplementary Figure 3. Antibody fusion proteins for delivery of soluble TNFLs.**

a, SEC chromatograms from preparative purification of trastuzumab and TNFα–trastuzumab fusion proteins (HC_mTNFα, LC_mTNFα and HCLC_mTNFα). Main peaks were collected for downstream analyses. Under these conditions, all fusion proteins eluted earlier than trastuzumab. b, SDS–PAGE analysis of purified samples under non-reducing and reducing conditions. Banding patterns are consistent with correct heavy- and light-chain assembly for each construct.

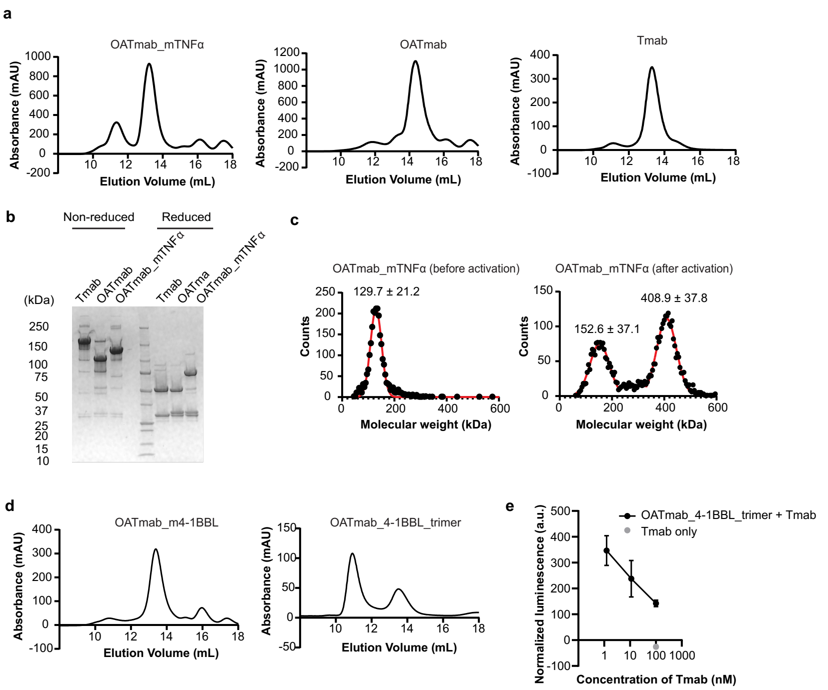


**Supplementary Figure 4. Antibody fusion proteins for delivery of membrane-type TNFLs.**

a, SEC chromatograms from preparative purification of the masked TNFα–trastuzumab fusion protein (OATmab_mTNFα) and control antibody formats (OATmab and Tmab). Main peaks were collected for downstream analyses. b, SDS–PAGE analysis of purified samples under non-reducing and reducing conditions. Banding patterns are consistent with correct heavy- and light-chain assembly for each construct. c, Mass photometry analysis of OATmab_mTNFα before (left) and after (right) TEV treatment. The shift in the mass distribution is consistent with TEV-dependent antibody homotrimerization. d, SEC chromatograms from preparative purification of the masked 4-1BBL–trastuzumab fusion protein (OATmab_m4-1BBL) and the homotrimeric fusion protein (OATmab_4-1BBL_trimer). Main peaks were collected for downstream analyses. e, Competition assay under the SK-BR-3 co-culture condition. Trimeric antibody (2 nM) was tested in the presence of increasing concentrations of Tmab. Data points show the mean of two technical replicates, and error bars indicate the individual replicate values (n = 2).

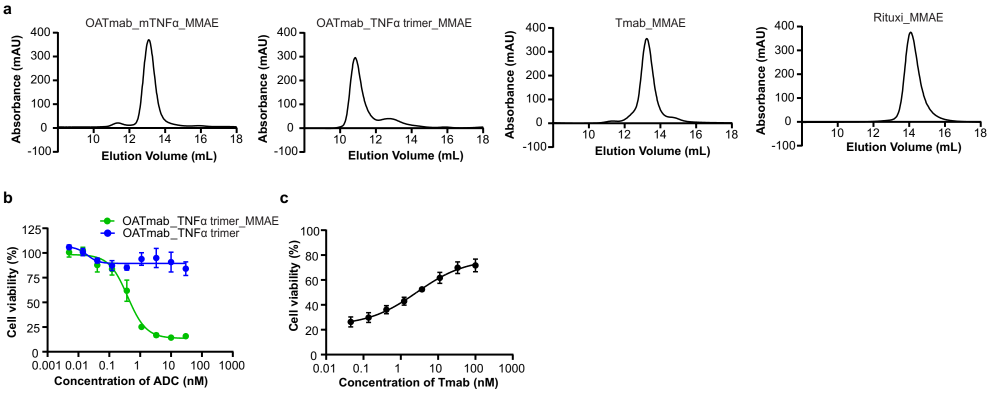


**Supplementary Figure 5. Purification of antibody–drug conjugates (ADCs) and cytotoxic activity.**

a, SEC chromatograms from preparative purification of ADCs. Main peaks were collected for downstream analyses. b, Cytotoxic activity of trimeric ADC (green) and unconjugated trimeric protein (blue) in HER2-positive SK-BR-3 cells. c, Competition assay in SK-BR-3 cells. Trimeric ADC (1 nM) was tested in the presence of increasing concentrations of unconjugated Tmab. Data are shown as mean ± s.d. from n = 3 technical replicates.

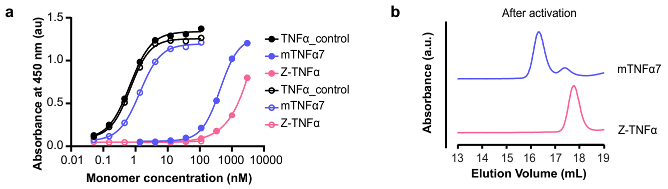


**Supplementary Figure 6. Binding and oligomeric-state analysis of masked TNFα using a previously reported Z-domain as a masking module.**

a, TNFR2 binding of masked TNFα constructs measured by ELISA before (filled circles) and after (open circles) TEV activation. Data are from a single experiment and are shown as representative results. Z-TNFα, which uses a previously reported Z-domain as the masking module, showed no detectable recovery of TNFR2 binding after TEV activation, whereas the selected de novo design mTNFα7 showed increased binding upon TEV treatment. b, Oligomeric states after TEV treatment assessed by analytical SEC. Z-TNFα eluted later than mTNFα7, which formed a trimer-like species after activation, indicating that Z-TNFα did not assemble into a trimeric TNFα species under these conditions. Linker cleavage was confirmed by SDS–PAGE (data not shown).

**Supplementary Table 1. AF2 metrics of masked TNFα**

| Protein | pLDDT | iPTM | RMSD (Å) | Masking effect | Characterization |
|---|---|---|---|---|---|
| TNFα_control | | | | | |
| mTNFα1 | 88.58 | 0.92 | 0.90 | Yes | |
| mTNFα2 | 88.00 | 0.92 | 0.91 | Yes | |
| mTNFα3 | 89.55 | 0.91 | 1.17 | Yes | Yes |
| mTNFα4 | 87.57 | 0.91 | 1.01 | Yes | Yes |
| mTNFα5 | 88.55 | 0.91 | 1.05 | | |
| mTNFα6 | 88.27 | 0.91 | 0.99 | Yes | |
| mTNFα7 | 88.12 | 0.90 | 0.96 | Yes | Yes |
| mTNFα8 | 88.61 | 0.90 | 0.92 | Yes | Yes |
| mTNFα9 | 89.06 | 0.89 | 0.94 | Yes | |
| mTNFα10 | 88.28 | 0.89 | 1.17 | | |
| mTNFα11 | 89.26 | 0.89 | 1.32 | | |
| mTNFα12 | 87.21 | 0.89 | 0.89 | | |
| mTNFα13 | 88.09 | 0.89 | 0.93 | | |
| mTNFα14 | 89.63 | 0.89 | 0.88 | | |

| | | | | |
|---|---|---|---|---|
| mTNFα15 | 88.45 | 0.88 | 0.99 | |
| mTNFα16 | 87.88 | 0.88 | 1.09 | |
| mTNFα17 | 87.76 | 0.88 | 0.91 | |
| mTNFα18 | 87.67 | 0.87 | 1.11 | |
| mTNFα19 | 88.40 | 0.86 | 1.28 | |
| mTNFα20 | 87.73 | 0.86 | 0.89 | |
| mTNFα21 | 89.52 | 0.90 | 0.98 | Yes |

**Supplementary Table 2. EC50 value of binding to TNFR2 before and after activation**

| Protein | EC50 | | |
|---|---|---|---|
| | Before activation (nM) | After activation (nM) | Ratio (Before/After) |
| TNFα_control | 0.67 (n = 1) | 0.69 (n = 1) | 0.97 |
| mTNFα3 | 256.7 (n = 1) | 30.4 (n = 1) | 8.4 |
| mTNFα4 | 185.4 (n = 1) | 1.22 (n = 1) | 152 |
| mTNFα7 | 431.2 (n = 1) | 1.37 (n = 1) | 315 |
| mTNFα8 | 572.4 (n = 1) | 2.95 (n = 1) | 194 |

**Supplementary Table 3. EC50 value of binding to Adalimumab before and after activation**

| Protein | EC50 | | |
|---|---|---|---|
| | Before activation (nM) | After activation (nM) | Ratio (Before/After) |
| TNFα_control | 0.54 (n = 1) | 0.75 (n = 1) | 0.72 |
| mTNFα3 | 590.8 (n = 1) | 36.2 (n = 1) | 16.3 |
| mTNFα4 | 211.5 (n = 1) | 1.15 (n = 1) | 184 |
| mTNFα7 | 752.2 (n = 1) | 1.42 (n = 1) | 530 |
| mTNFα8 | 556.5 (n = 1) | 4.26 (n = 1) | 131 |

**Supplementary Table 4. EC50 value of biological function to WEHI-13VAR before and after activation**

| Protein | EC50 | | |
| --- | --- | --- | --- |
| | Before activation (pM) | After activation (pM) | Ratio (Before/After) |
| TNFα_control | 1.87 (n = 3) | 1.67 (n = 3) | 1.1 |
| mTNFα3 | 2450 (n = 3) | 583 (n = 3) | 4.2 |
| mTNFα4 | 148 (n = 3) | 2.30 (n = 3) | 64 |
| mTNFα7 | 184 (n = 3) | 2.49 (n = 3) | 74 |
| mTNFα8 | 517 (n = 3) | 11.7 (n = 3) | 44 |
| Trastuzumab | | | |
| HC fusion | 26.2 (n = 3) | 1.16 (n = 3) | 23 |
| LC fusion | 46.0(n = 3) | 1.05 (n = 3) | 44 |
| HCLC fusion | 19.1 (n = 3) | 0.66 (n = 3) | 29 |

**Supplementary Table 5. AF2 metrics of masked OX40L**

| Protein | Design method | Template | pLDDT | iPTM | RMSD (Å) | Masking effect | Characterization |
|---|---|---|---|---|---|---|---|
| OX40L_control | | | | | | | |
| mOX40L1 | 1 | | 89.46 | 0.89 | 0.52 | Yes | Yes |
| mOX40L2 | 1 | | 90.59 | 0.89 | 0.79 | Yes | |
| mOX40L3 | 1 | | 89.56 | 0.89 | 0.72 | | |
| mOX40L4 | 2 | mTNFα7 | 91.53 | 0.91 | 0.67 | Yes | Yes |
| mOX40L5 | 2 | mTNFα7 | 90.06 | 0.89 | 0.75 | Yes | |
| mOX40L6 | 2 | mTNFα7 | 90.47 | 0.89 | 0.71 | Yes | |
| mOX40L7 | 1 | | 90.27 | 0.89 | 0.78 | Yes | Yes |
| mOX40L8 | 2 | mTNFα7 | 90.12 | 0.89 | 0.77 | Yes | Yes |
| mOX40L9 | 2 | mTNFα7 | 91.10 | 0.89 | 0.64 | Yes | |
| mOX40L10 | 1 | | 90.96 | 0.89 | 0.84 | | |
| mOX40L11 | 1 | | 90.99 | 0.88 | 0.78 | Yes | |
| mOX40L12 | 1 | | 90.91 | 0.91 | 0.34 | Yes | |
| mOX40L13 | 1 | | 91.69 | 0.91 | 0.43 | Yes | |
| mOX40L14 | 1 | | 91.00 | 0.90 | 0.40 | Yes | |

| | | | | | |
|---|---|---|---|---|---|
| mOX40L15 | 1 | 91.36 | 0.90 | 0.40 | Yes |
| mOX40L16 | 1 | 90.50 | 0.90 | 0.38 | Yes |
| mOX40L17 | 1 | 90.56 | 0.90 | 0.61 | Yes |
| mOX40L18 | 1 | 90.26 | 0.90 | 0.41 | Yes |

**Supplementary Table 6. AF2 metrics of masked 4-1BBL**

| Protein | Design method | Template | pLDDT | iPTM | RMSD (Å) | Masking effect | Characterization |
|---|---|---|---|---|---|---|---|
| 4-1BBL_control | | | | | | | |
| m4-1BBL1 | 1 | | 92.73 | 0.89 | 0.73 | Yes | |
| m4-1BBL2 | 1 | | 92.27 | 0.88 | 0.54 | Yes | Yes |
| m4-1BBL3 | 1 | | 92.21 | 0.90 | 1.30 | Yes | |
| m4-1BBL4 | 2 | mTNFα8 | 92.93 | 0.90 | 0.84 | Yes | |
| m4-1BBL5 | 2 | mTNFα8 | 93.89 | 0.90 | 0.71 | Yes | Yes |
| m4-1BBL6 | 2 | mTNFα8 | 93.58 | 0.90 | 0.64 | Yes | Yes |
| m4-1BBL7 | 1 | | 92.11 | 0.88 | 0.66 | Yes | |
| m4-1BBL8 | 2 | mTNFα7 | 93.15 | 0.90 | 1.37 | Yes | Yes |
| m4-1BBL9 | 1 | | 92.85 | 0.90 | 0.54 | Yes | |
| m4-1BBL10 | 1 | | 93.00 | 0.89 | 0.69 | Yes | |
| m4-1BBL11 | 2 | mTNFα7 | 91.75 | 0.89 | 0.57 | Yes | |
| m4-1BBL12 | 1 | | 92.27 | 0.89 | 0.75 | Yes | |
| m4-1BBL13 | 3 | mTNFα8 | 93.18 | 0.88 | 0.58 | Yes | Yes |
| m4-1BBL14 | 1 | | 93.05 | 0.88 | 0.87 | Yes | |

| | | | | | | |
|---|---|---|---|---|---|---|
| m4-1BBL15 | 1 | | 92.65 | 0.88 | 0.61 | Yes |
| m4-1BBL16 | 3 | mTNFα8 | 93.06 | 0.88 | 0.73 | Yes |
| m4-1BBL17 | 3 | mTNFα8 | 91.85 | 0.88 | 0.58 | Yes |

**Supplementary Table 7. EC50 value of binding to OX40 before and after activation**

| Protein | EC50 | | |
|---|---|---|---|
| | Before activation (nM) | After activation (nM) | Ratio (Before/After) |
| OX40L_control | 0.60 (n = 1) | 0.68 (n = 1) | 0.88 |
| mOX40L1 | 344.1 (n = 1) | 0.98 (n = 1) | 351 |
| mOX40L4 | >1,500 (n = 1) | 1.27 (n = 1) | >1,000 |
| mOX40L7 | 273.0 (n = 1) | 0.81 (n = 1) | 337 |
| mOX40L8 | >1,500 (n = 1) | 1.23 (n = 1) | >1,000 |

**Supplementary Table 8. EC50 value of binding to TNFR2 before and after activation**

| Protein | EC50 | | |
|---|---|---|---|
| | Before activation (nM) | After activation (nM) | Ratio (Before/After) |
| Trastuzumab | | | |
| HC fusion | 64.1 (n = 1) | 0.76 (n = 1) | 84.3 |
| LC fusion | 58.1 (n = 1) | 0.67 (n = 1) | 86.7 |
| HCLC fusion | 5.32 (n = 1) | 0.18 (n = 1) | 29.6 |

**Supplementary Table 9. EC50 value of binding to HER2**

| Protein | EC50 (nM) |
| --- | --- |
| Trastuzumab | 0.23 (n = 1) |
| HC fusion | 0.37 (n = 1) |
| LC fusion | 0.28 (n = 1) |
| HCLC fusion | 0.50 (n = 1) |

**Supplementary Table 10. EC50 value of binding to HER2**

| Protein | EC50 (nM) |
|---|---|
| Tmab | 0.068 (n = 3) |
| OATmab_mTNFα | 1.49 (n = 3) |
| OATmab_TNFα trimer | 0.074 (n = 3) |

**Supplementary Table 11. EC50 value of binding to TNFR2**

| Protein | EC50 (nM) |
| --- | --- |
| OATmab_mTNFα | 6.95 (n = 3) |
| OATmab_TNFα trimer | 0.031 (n = 3) |

**Supplementary Table 12. IC50 value of cytotoxic activity of ADCs to SK-BR-3 cell**

| ADC | IC50 (nM) |
|---|---|
| OATmab_mTNFα_MMAE | 12.5 (n = 3) |
| OATmab_TNFα trimer_MMAE | 0.39 (n = 3) |
| Tmab_MMAE | 0.32 (n = 3) |
| Rituximab_MMAE | 28.0 (n = 3) |
